# Molecular Characterization of Carbapenem-Resistant *Acinetobacter baumannii* Clinical Isolates from Egyptian Patients

**DOI:** 10.1101/2020.08.24.264911

**Authors:** Reem M Hassan, Sherifa T Salem, Saly Ismail Mostafa Hassan, Asmaa Sayed Hegab, Yasmine S Elkholy

**Affiliations:** Department of Clinical and Chemical Pathology, Faculty of Medicine, Cairo University, Egypt; Department of Clinical Pathology, Faculty of Medicine, Beni-Suef University, Egypt; Department of Medical Microbiology and Immunology, Faculty of Medicine, Cairo University, Egypt

## Abstract

*Acinetobacter baumannii* (*A. baumannii*) represents a global threat owing to its ability to resist most of the currently available antimicrobial agents. Moreover, emergence of carbapenem resistant *A. baumannii* (CR-AB) isolates limits the available treatment options. Enzymatic degradation by variety of ß-lactamases, have been identified as the most common mechanism of carbapenem resistance in *A. baumannii*. The alarming increase in the prevalence of CR-AB necessitates continuous screening and molecular characterization to appreciate the problem. The present study was performed to assess the prevalence and characterize carbapenemases among 206 CR-AB isolated from various clinical specimens collected from different intensive care units at Kasr Al-Aini Hospital.

All isolates were confirmed to be *A. baumannii* by detection of the blaOXA-51-like gene. Molecular screening of 13 common Ambler class bla carbapenemases genes in addition to insertion sequence (IS-1) *upstream* OXA-23 was performed by using four sets of multiplex PCR, followed by identification using gene sequencing technology. Among the investigated genes, the prevalence of blaOXA-23, and blaOXA-58 were 77.7%, and 1.9%, respectively. The IS*Aba1* was detected in 10% of the blaOXA-23 positive isolates. The prevalence of metallo-β-lactamases (MBLs) studied; blaNDM-1, blaSPM, blaVIM, blaSIM-1 were 11.7%, 6.3%, 0.5%, and 0.5% respectively. One of class A; bla KPC was detected in 10.7% of the investigated isolates. blaOXA-24/40, blaIMP, blaGES, blaVEB and blaGIM were not detected in any of the studied isolates. Moreover, 18.4% of the isolates have shown to harbor two or more of the screened bla genes. We concluded that the most prevalent type of ß-lactamases genes among CR-AB isolates collected from Egyptian patients were blaOXA-23 followed by blaNDM-1 and blaKPC.

**Author summary:** Carbapenem-resistant *A. baumannii* has become a real global health threat. The aim of the present study was to characterize and to assess the prevalence of carbapenemases among 206 CR-AB clinical isolates from Egyptian patients. We concluded that the most prevalent type of ß-lactamases genes among CR-AB isolates collected from Egyptian patients were blaOXA-23 followed by blaNDM-1 and blaKPC. In this study, IS*Aba1* was detected upstream 10% of blaOXA-23 positive isolates only which indicates that the spread of resistance among *Acinetobacter* isolates could be either chromosomal or plamid-mediated.

## Introduction

*A. baumannii* has been identified as an opportunistic nosocomial pathogen, which is tolerant to wide ranges of temperature, pH, and humidity and is able to survive on dry surfaces for months, so it is recognized as a real challenge to infection control measures (1). It causes a wide variety of infections including pneumonia, urinary tract infection, skin and soft tissue infections, central line associated bloodstream infections and others (2).

*A. baumannii* represents a global threat owing to its ability to resist most of the currently available antimicrobial agents including ß-lactams, fluoroquinolones and aminoglycosides (3). Un-regulated use of antibiotics in healthcare settings results in emergence of multidrug-resistance added to the intrinsic resistance of *A. baumannii* especially in intensive care units (4). Moreover, emergence of carbapenem resistant *A. baumannii* isolates limit the available treatment options for such infections (5).

The most common mechanism responsible for carbapenem resistance among *A. bauminii* is enzymatic degradation by variety of ß-lactamases, the four β-lactamase classes (A, B, C and D) have been detected in *A. baumannii* (6, 7, 8)

Two intrinsic types of ß-lactamases can be identified in almost all *A. baumannii* isolates; AmpC-type cephalosporinases and OXA-51/69 variants. Both of which are chromosomally located, and have little impact on carbapenems, if any (9).

However, higher carbapenem hydrolysis rates have been reported to result from the acquisition of insertion sequences (ISs) that affect the expression of blaOXA genes encoding oxacillinases (10).These elements are the most abundant transposable elements capable of causing mutations and rearrangements in the genome, contributing to the spread of resistance and virulence determinants among bacterial species (11). Insertion sequence *A. baumannii*-1 (IS*Aba1)* upstream blaOXA-51 like genes, belong to the IS4 family and has been associated with increased gene expression and hence carbapenem resistance (8,12).

Till date, carbapenem-hydrolyzing class D β-lactamases, the so called (CHDLs) also named oxacillinases (OXAs) for their effect on oxacillin (13) seem to be the most common mechanism of carbapenem resistance in *A. baumannii* (8). Class D β-lactamases, includes six subgroups: The intrinsic OXA-51 and the acquired OXA-23-like, OXA-58-like, OXA-24/40-like, OXA-235-like and OXA-143-like β-lactamases (13).

Additionally, class B β-lactamases, also known as MBLs play a less important yet more potent role in carbapenem resistance among *A. baumannii* isolates (14).

Four MBLs are known in *A. baumannii*: New Delhi metallo-β-lactamase (NDM), Imipenemase (IMP), Seoul Imipenemase (SIM) and Verona integron-encoded metallo-β-lactamase (VIM) (14, 15). Two variants of NDM (NDM-1 and NDM-2) have been reported in *A. baumannii* clinical isolates in Egypt (16, 17).

On the contrary of class B and D, few of class A including *K. pneumonia* carbapenemase (KPC) type enzymes; blaKPC and blaGES were also incriminated of carbapenem resistance among *A. baumannii* (18).

Other ESBLs were identified among *A*.*baumannii* isolates and confer resistance to broad spectrum cephalosporins e.g. blaPER, blaVEB and others (19).

Despite of the global increase at alarming rates of CR-AB, few studies were devoted to this organism in Egypt (20, 21, 22). More data are crucial to appreciate the problem objectively and to control further evolution of CR-AB.

The aim of the present study was to characterize and to assess the prevalence of carbapenemases among 206 CR-AB clinical isolates from Egyptian patients.

## Results

A total of 206 clinical isolates of CR-AB were collected from different clinical specimens mainly from different ICUs (n=154, 74.8%) at Kasr Al-Aini Hospital. Most of the CR-AB detected were isolated from wound swabs (n=77), respiratory secretions (n=56), blood cultures (n=37), and urine samples (n=27) **(Table 1)**.

**Table 1:** Distribution of *A. baumannii* isolates among different clinical specimens.

| Clinical Specimen | *A. baumannii* isolates<br>N (%) |  |
| --- | --- | --- |
| Wound discharge | 77 | (37.4%) |
| Respiratory secretions | 56 | (27.2%) |
| Blood | 37 | (18.0%) |
| Urine | 27 | (13.1%) |
| Others (body fluids, drains) | 9 | (4.4%) |

All isolates were confirmed to be *A. bauminii* by detecting blaOXA-51-like gene among all (100%) the studied isolates **(Table2)**.

Among the investigated genes, class D blaOXA-23 was the most commonly detected gene (77.7%) and the *ISAba1*was detected only in 10% of the blaOXA-23 positive isolates.

One of the MBLs; blaNDM was detected in 11.7%, and one of class A; blaKPC was detected in 10.7% of the investigated isolates. Other carbapenemases were detected at lower frequencies. However, five of the investigated genes (blaOXA-24, blaIMP, blaGES, blaVEB and blaGIM)were not detected in any of the studied isolates **(Table 2)**.

**Table 2:** Distribution of different bla gene classes and genes among the CR-AB isolates.

| bla class | bla gene | *A. baumannii* isolates<br>N (%) |  |
| --- | --- | --- | --- |
| Class A | blaKPC | 22 | (10.7%) |
|  | blaGES | 0 | (0.0%) |
| Class-B | blaVIM | 1 | (0.5%) |
|  | blaIMP | 0 | (0.0%) |
|  | blaGIM | 0 | (0.0%) |
|  | blaSPM | 13 | (6.3%) |
|  | blaSIM-1 | 1 | (0.5%) |
|  | blaNDM-1 | 24 | (11.7%) |
| **Class D** | blaOXA-51 | 206 | (100%) |
|  | blaOXA-23 | 160 | (77.7%) |
|  | blaOXA-24/40 | 0 | (0.0%) |
|  | blaOXA-58 | 4 | (1.9%) |
| **Other beta-lactamases** | blaPER | 2 | (1.0%) |
|  | blaVEB | 0 | (0.0%) |
| **ISAb1 upstream of**<br>**blaOXA-23** |  | 16 | (7.8%) |

**Table 3:**
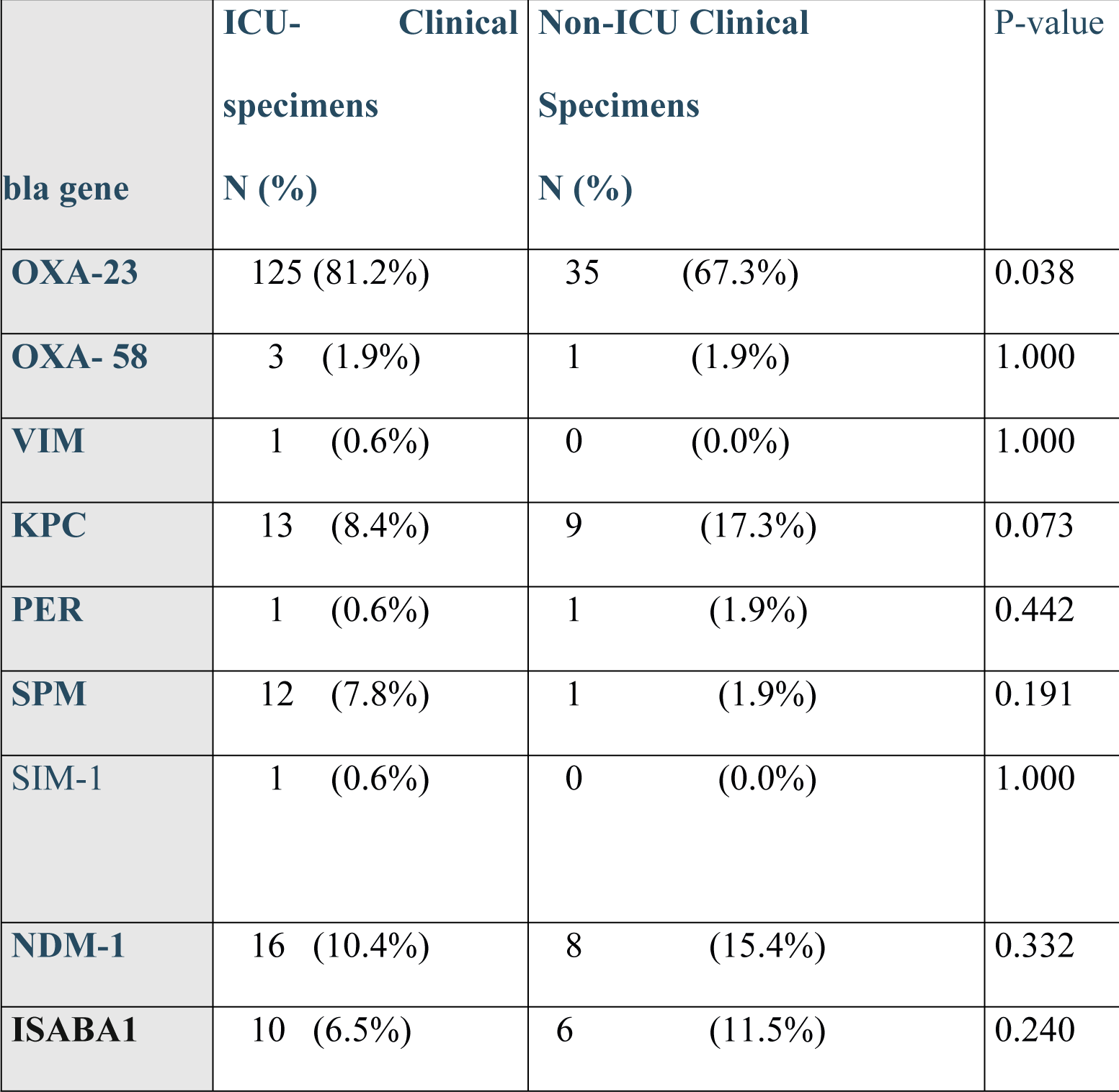
Difference in distribution of bla genes among CR-AB isolates from ICU and Non-ICU clinical specimens.

**Non-ICU clinical specimens**
| **bla gene** | **ICU- Clinical specimens**<br>**N (%)** | **Non-ICU Clinical Specimens**<br>**N (%)** | **P-value** |
| --- | --- | --- | --- |
| **OXA-23** | 125 (81.2%) | 35 (67.3%) | 0.038 |
| **OXA- 58** | 3 (1.9%) | 1 (1.9%) | 1.000 |
| **VIM** | 1 (0.6%) | 0 (0.0%) | 1.000 |
| **KPC** | 13 (8.4%) | 9 (17.3%) | 0.073 |
| **PER** | 1 (0.6%) | 1 (1.9%) | 0.442 |
| **SPM** | 12 (7.8%) | 1 (1.9%) | 0.191 |
| **SIM-1** | 1 (0.6%) | 0 (0.0%) | 1.000 |
| **NDM-1** | 16 (10.4%) | 8 (15.4%) | 0.332 |
| **ISABA1** | 10 (6.5%) | 6 (11.5%) | 0.240 |

Although most of the studied CR-AB isolates were detected in specimens from different ICUs, there was no significant difference in the prevalence of the investigated genes in clinical specimens collected from the ICUs and specimens collected from other departments **(Table3)**.

Moreover, 18.4% of the isolates (n=38) have shown to harbor two or more of the tested bla genes **(Table 4)**.

**Table 4:** CR-AB isolates harbor more than one of the bla genes:

| **bla genes** | **CR-AB isolates harbor more than one *bla* genes (N=38)** |
| --- | --- |
| OXA-23, KPC | 10 |
| OXA-23, NDM | 6 |
| OXA-23, SPM | 6 |
| OXA23, KPC, NDM | 6 |
| OXA-58, NDM | 3 |
| OXA-23, PER | 2 |
| OXA-23, NDM, SPM | 2 |
| OXA-23, SIM | 1 |
| OXA-23, OXA-58, KPC | 1 |
| OXA-23, NDM, VIM | 1 |

## Materials and Methods

This cross sectional study involved a total of 206 clinical, non-duplicate, isolates of CR-AB.; which were collected over one year duration from December 2018 to December 2019. The research study was approved by the institutional Review Board of Kasr Al-Aini Hospital and the Research Ethics Committee at Faculty of Medicine, Cairo University.

### I. Bacterial Isolates

The isolates were collected at the central microbiology laboratory at Kasr Al-Aini Hospital. The isolates were obtained from different clinical specimens, including wound swabs, respiratory secretions, blood cultures, urine samples and others including body fluid and drains collected from inpatients admitted at intensive care units and other different departments at Kasr Al-Aini Hospital. The isolates were identified phenotypically by colonial morphology, Gram-staining and conventional biochemical testing. All isolates grown as lactose non fermenter colonies on MacConkey agar, showing Gram negative coccobaciili were preliminary identified as *Acinetobater* spp. (23).

### II. Carbapenem susceptibility testing

Demonstration of carbapenem susceptibility among all of the initially identified *Acinetobacter* clinical isolates was conducted by the standard disc diffusion technique on Müller-Hinton agar using imipenem and meropenem discs (Oxoid, Basingstoke, United Kingdom), and interpreted following the Clinical and Laboratory Standards Institute (CLSI) (24).

### III. Molecular Identification

All CR-AB isolates were selected and submitted to DNA extraction by the heat shock method, followed by genotypic identification of *A. baumannii* by detection of the blaOXA-51-like gene (25). The 206 isolates were confirmed to be *A. baumannii* were further investigated.

### IV. Molecular detection of different carbapenemases encoding genes and insertion sequence (IS) blaOXA-23

Four sets of multiplex PCR were done including common Ambler class bla genes that cause carbapenem resistance; multiplex 1 included blaOXA-23, blaOXA-24 and blaOXA-58 (26,27), multiplex 2 included blaVIM, blaKPC and blaIMP, multiplex 3 included blaGES, blaPER and blaVEB (28), while multiplex 4 contained blaGIM, blaSIM-1, blaSPM and blaNDM-1 (28,29). Presence of IS*Aba1* upstream of blaOXA-23 was investigated using IS*Aba1*F/OXA-23-likeR (12). PCR was performed using PCR-EZ D-PCR Master Mix (Bio Basic Inc., Canada) in a Bio-Rad Thermal Cycler PTC-200. Briefly, an initial denaturation step of 95 °C for 15 min, followed by 30 cycles of denaturation at 94 °C for 30 sec, an annealing temperature dependent on the melting temperature of the primer pair (multiplex I and IS*Aba1*F/OXA-23-likeR: 52 °C; multiplex II and III: 57 °C and multiplex IV: 60 °C) and extension at 72 °C for 90 sec, followed by the final extension step at 70 °C for 10 min. A negative control (sterile nuclease free water) was included for all PCR assays. Amplified PCR products were purified using PureLink® PCR Purification Kit (Invitrogen, Carlsbad, CA, USA) according to manufacturer’s instructions. A BigDye Terminator v3.1 cycle sequencing kit (Applied Biosystems, Foster City, CA, USA) was used to sequence the PCR amplified products from the positive cases according to manufacturer’s instructions. The sequenced products were run on a 3500 Genetic Analyzer (Applied Biosystems). Sequences were compared with those available in the GenBank database using the basic local alignment search tool (BLAST, www.ncbi.nlm.nih.gov).

## Discussion

Outlining the genetic background of carbapenem resistance among 206 *A. baumannii* isolates collected from Kasr Al Aini Hospital, one of the largest tertiary care hospitals in Egypt, was the main scope of the present study. As proved by other studies, bla OXA-51, is an oxacillinase present in all *A. baumannii* isolates. Luckily, it is too weak to confer by itself carbapenem resistance. However, it is frequently used to confirm the identity of *A*.*baumannii* (27, 30). All the investigated isolates of our study were confirmed to harbor this gene.

In concordance with many other studies, the most common carbapenemase gene was found to be blaOXA-23 that belongs to Class D β-lactamases. In the present study, it was detected in 77.7% of the investigated isolates. In Egypt, a study in Zagazig University Hospitals reported that 90% of 50 carbapenem resistant *A. baumannii* isolates were detected to harbor blaOXA-23 (31). Obviously the higher carriage rate in the later study may be attributed to the fact that all isolates were collected from surgical ICUs, whereas in our study the investigated isolates were isolated from both ICU and non ICU settings. Other studies in Egypt and Saudi Arabia showed higher prevalence rate (100%) of blaOXA-23 among carbapenem resistant *A. baumannii* isolates (32, 33). IS*Aba1* was detected upstream blaOXA-23 gene only in 10% of the blaOXA-23 positive isolates. These findings confirmed that the blaOXA-23 gene could be located on the chromosome or on a plasmid (34). We can conclude that the current worldwide dissemination of the blaOXA-23 gene is associated with different genetic structures and plasmids. The dynamic spread of blaOXA-23 will make it difficult to control because this spread is not associated with a single entity (35).

Within the same class; class D β-lactamases, blaOXA-58 was detected only among 1.9% of our studied isolates. Closely related to our results, other studies in Egypt, in USA, and in Palestine could detect blaOXA-58 among 1.4%, 2%, and 3% respectively among carbapenem resistant *A. baumannii* investigated isolates (22, 36, 37). However, another Egyptian study couldn’t find the former gene among any of their studied carbapenem resistant *A. baumannii* isolates (38). Although, a higher blaOXA-58 prevalence was reported among carbapenem resistant *A. baumannii* isolates in Egypt (9.1%) (39) and Algeria (14.7%) (40). Therefore, more studies are needed from different region in Egypt and moreover from different countries before we can conclude that the prevalence of blaOXA-58 is low among carbapenem resistant *A. baumannii*.

Till date the most common carbapenemsases are carbapenem-hydrolyzing class D β-lactamases (CHDLs) and, to a lesser extent, MBLs. Although MBLs has been reported in sporadic parts of the world (41), most of MBLs were first described in *A. baumannii* in Egypt (16, 17, 42) then emerged in the middle east (43). In the present study MBLs were detected in 19% of the investigated isolates; with blaNDM accounts for 11.7%, blaSPM 6.3%, blaVIM 0.5%, and blaSIM 0.5%. Nevertheless, in the present study, none of the *A. baumannii* isolates were found to harbor blaIMP, nor blaGIM.

In Egypt many authors have reported different prevalence of MBLs in *A. baumannii. Elkasaby and El Sayed Zaki, (2017*) reported that among 280 *A. baumannii* isolates collected from Egyptian patients admitted to Mansoura University Hospital ICU, 95% harbored MBLs, of which, blaIMP accounted for 95.7%. However, the authors didn’t investigate blaNDM among the studied isolates (21). Another Egyptian study has detected blaNDM among 66.7% of 50 *A*.*baumannii* isolates. Yet, the former study didn’t find neither blaIPM nor blaVIM among the tested isolates (31). Another study in Egypt has shown that blaNDM accounted for carbapenem resistance in 27.58%, and blaVIM in 10.3% of studied isolates (20). Two MBLs: blaVIM and blaNDM were detected in 100% and 12.1% respectively in seventy four CR-AB investigated isolates, collected from different clinical specimens at Alexandria University Hospital (32). In Palestine Handal and colleagues reported that *bla*NDM was detected among 5.8% of 69 carbapenem resistant *A. baumannii* isolates (37).

In the present study, two of the Class A carbapenemases, namely blaKPC and blaGES were tested. While none of isolates were shown to harbor blaGES, blaKPC was detected in 10.7% of the investigated isolates. Closely related results were detected in other Egyptian studies. bla KPC were not detected among 40 carbapenem resistant *A. baumannii* isolates collected from 2 hospitals in Egypt (22, 31, 32), while blaGES was detected among 50% of the investigated isolates in the later same study (32).

## Conclusion

We can conclude that the worldwide spread of carbapenem-resistant *A. baumannii* has become a real global health threat. Among the investigated isolates in the present study CR-AB were confirmed by detection of blaOXA-51. Class D carbapenemase blaOXA-23 was the most prevalent followed by blaNDM-1 belonging to class B MBLs and class A blaKPC. In this study, IS*Aba1* was detected upstream 10% of blaOXA-23 positive isolates only which indicates that the spread of resistance among *Acinetobacter* isolates could be either chromosomal or plamid-mediated. Further investigations should be continue to appreciate the reality of the problem of multi-drug resistant *A. baumannii*.

## References

1. Kramer A, Schwebke I, Kampf G. How long do nosocomial pathogens persist on inanimate surfaces? A systematic review. BMC Infectious Diseases. 2006; 6(130).

2. Cai Y, Chai D, Wang R, Liang B, Bai N. Colistin resistance of *Acinetobacter baumannii*: clinical reports, mechanisms and antimicrobial strategies. Journal of Antimicrobial Chemotherapy. 2012; 67(7):1607–15.

3. Peleg AY, Seifert H, Paterson DL. *Acinetobacter baumannii*: emergence of a successful pathogen. Clinical Microbiology Reviews. 2008; 21(3):538–82.

4. Almasaudi SB. *Acinetobacter* spp. as nosocomial pathogens: epidemiology and resistance features. Saudi J Biol Sci. 2018; 25(3):586–96.

5. Fattouh M, El-din AN. Emergence of carbapenem resistant *Acinetobacter baumannii* in the intensive care unit in Sohag University hospital, Egypt. Int J Curr Microbiol App Sci. 2014; 3(4):732–44.

6. Paton R, Miles RS, Hood J, Amyes SG. ARI 1: Beta-lactamase-mediated imipenem resistance in *Acinetobacter baumannii*. Int J Antimicrob Agents. 1993, Feb; 2 (2):81–7.

7. Scaife W, Young HK, Paton RH, Amyes SG. Transferable imipenem-resistance in *Acinetobacter* species from a clinical source. J Antimicrob Chemother. 1995; 36(3):585–6.

8. Poirel L, Nordmann P. Carbapenem resistance in *Acinetobacter baumannii*: mechanisms and epidemiology. Clin Microbiol Infect. 2006; 12:826–836.

9. Héritier C, Poirel L, Fournier PE, Claverie JM, Raoult D, Nordmann P. Characterization of the naturally occurring oxacillinase of *Acinetobacter baumannii*. Antimicrob.Agents Chemother. 2005; 49(10), 4174–79.

10. Peleg AY, Seifert H, Paterson DL. *Acinetobacter baumannii*: emergence of a successful pathogen. Clinical Microbiology Reviews. 2008; 21(3):538–82.

11. Vandecraena J, Chandlerc M, Aertsenb A, Houdta RV. The impact of insertion sequences on bacterial genome plasticity and adaptability. Critical Reviewes in Microbiology. 2017; 43 (6): 709–30.

12. Turton JF, Ward ME, Woodford N, Kaufmann ME, Pike R, Livermore DM, Pitt TL. The role of IS*Aba1* in expression of OXA carbapenemase genes in *Acinetobacter baumannii*. FEMS Microbiol. Lett. 2006; 258(1), 72–7.

13. Poirel L, Naas T, Nordmann P. Diversity, epidemiology, and genetics of class D β-lactamases. Antimicrob Agents Chemother. 2010: 54(1):24–38.

14. Walsh TR, Toleman MA, Poirel L, Nordmann P. Metallo-β-lactamases: the quiet before the storm? Clin Microbiol Rev 2005; 18: 306–25.

15. Bonnin R, Poirel L, Naas T, Pirs M, Seme K, Schrenzel J, Nordmann p. Dissemination of New Delhi metallo-β-lactamase-1-producing *Acinetobacter baumannii* in Europe. Clin Microbiol Infect. 2012; 18(9): 362–5.

16. Hrabák J, Štolbová M, Študentová V, Fridrichová M, Chudáčková E, Zemlickova H. NDM-1 producing *Acinetobacter baumannii* isolated from a patient repatriated to the Czech Republic from Egypt, July 2011. Eurosurveillance. 2012; 17(7):1–3.

17. Kaase M, Nordmann P, Wichelhaus TA, Gatermann SG, Bonnin RA, Poirel L. NDM-2 carbapenemase in *Acinetobacter baumannii* from Egypt. J Antimicrob Chemother. 2011; 66(6):1260–2.

18. Pogue JM, Mann T, Barber KE, Kaye KS. Carbapenem-resistant *Acinetobacter baumannii*: epidemiology, surveillance and management. Exp Rev Anti infect Ther. 2013; 11: 383–393.

19. Bonnin RA, Potron A, Poirel L, Lecuyer H, Neri R, Nordmann P. PER-7, an extended-spectrum beta-lactamase with increased activity toward broad-spectrum cephalosporins in *Acinetobacter baumannii*. Antimicrob Agents Chemother. 2011; 55:2424–2427.

20. Kamel NA, El-Tayeb WN, El-Ansary MR, Mansour MT, Aboshanab KM. Phenotypic screening and molecular characterization of carbapenemase-producing gram-negative bacilli recovered from febrile neutropenic pediatric cancer patients in Egypt. PLoS One 2018; 13(8).

21. Alkasaby NM, and El Sayed Zaki M. Molecular study of *Acinetobacter baumannii* isolates for metallo-beta-lactamases and extended-spectrum-betalactamases genes in intensive care unit, Mansoura university hospital, Egypt. Int. J. Microbiol. 2017; 3925868.

22. Al-Agamy MH, Khalaf NG, Tawfick MM, Shibl AM, Kholy AE. Molecular characterization of carbapenem-insensitive *Acinetobacter baumannii* in Egypt. Int J Infect Dis. 2014; 22:49–54.

23. Constantiniu S, Romaniuc A, Iancu LS, Filimon R, Taraşi I. Cultural and biochemical characteristics of *Acinetobacter* spp strains isolated from hospital units. J Prev Med. 2004; 12 (3–4):35–42.

24. Clinical and Laboratory Standards Institute Guidelines. M100-S28 performance standards for antimicrobial susceptibility testing. 2018

25. Turton JF, Woodford N, Glover J, Yarde S, Kaufmann ME, Pitt TL. Identification of *Acinetobacter baumannii* by detection of the blaOXA-51-like carbapenemase gene intrinsic to this Species. J Clin Microbiol. 2006; 44: 2974–2976.

26. Woodford N, Ellington MJ, Coelho JM, Turton JF, Ward ME, Brown S, Amyes SG, Livermore DM. Multiplex PCR for genes encoding prevalent OXA carbapenemases in *Acinetobacter* spp. Int J Antimicrob Agents. 2006; 27: 351–353.

27. Mostachio AK, Van Der Heidjen I, Rossi ., Levin AS, Costa SF. Multiplex PCR for rapid detection of genes encoding oxacillinases and metallo-β-lactamases in carbapenem resistant *Acinetobacter* spp. Journal of Medical Microbiology. 2009; 58(11):1522–1524.

28. Dallenne C, Da Costa A, Decré D, Favier C, Arlet G. Development of a set of multiplex PCR assays for the detection of genes encoding important β-lactamases in *Enterobacteriaceae*. J Antimicrob Chemother. 2010; 65: 490–495.

29. Voets GM, Fluit AC, Scharringa J, Stuart JC, Leverstein-van Hall MA. A set of multiplex PCRs for genotypic detection of extended-spectrum β-lactamases, carbapenemases, plasmid-mediated AmpC β-lactamases and OXA β-lactamases. Int J Antimicrob Agents. 2011; 37: 356–359.

30. Evans BA, Amyes SG. OXA β-lactamases. Clinical Microbiology Reviews. 2014; 27(2): 241–263.

31. Ramadan RA, Gebriel MG, Kadry HM, and Mosallem A. Carbapenem-resistant *Acinetobacter baumannii* and *Pseudomonas aeruginosa*: characterization of carbapenemase genes and E-test evaluation of colistin-based combinations. Infect. Drug Resist. 2018; 11: 1261–1269.

32. Abouelfetouh A, Torky AS, Aboulmagd E. Phenotypic and genotypic characterization of carbapenem-resistant *Acinetobacter baumannii* isolates from Egypt. Antimicrobial Resistance and Infection Control. 2019; 8:185.

33. El-Mahdy TS, Al-Agamy MH, Al-Qahtani AA, and Shibl AM. Detection of blaOXA-23-like and blaNDM-1 in *Acinetobacter baumannii* from the eastern region, Saudi Arabia. Microb. Drug Resist. 2014; 23: 115–121.

34. Kulkosky J, Jones KS, Katz RA, Mack JP, Skalka AM. Residues critical for retroviral integrative recombination in a region that is highly conserved among retroviral/retrotransposon integrases and bacterial insertion sequence transposases. Mol. Cell. Biol. 1992; 12: 2331–2338.

35. Mugnier PD, Poirel L, Nordmann P. Functional Analysis of Insertion Sequence ISAba1, Responsible for Genomic Plasticity of *Acinetobacter baumannii*. Journal of Bacteriology. 2009; 191 (7): 2414–18.

36. Huang XZ, Cash DM, Chahine MA, Nikolich MP, Craft DW. Development and validation of a multiplex TaqMan real-time PCR for rapid detection of genes encoding four types of class D carbapenemase in *Acinetobacter baumannii*. Journal of Medical Microbiology. 2012; 61: 1532–37.

37. Handal R, Qunibi L, Sahouri I, Juhari M, Dawodi R, Marzouqa H, Hindiyeh M. Characterization of Carbapenem-Resistant *Acinetobacter baumannii* Strains Isolated from Hospitalized Patients in Palestine. Int J Microbiol. 2017; 1–7.

38. Benmahmod AB, Said HS, Ibrahim RH. Prevalence and mechanisms of carbapenem resistance among *Acinetobacter baumannii* clinical isolates in Egypt. Microb Drug Resist. 2018; 25(4):480–8.

39. Al-Hassan L, El Mehallawy H, Amyes S. Diversity in *Acinetobacter baumannii* isolates from paediatric cancer patients in Egypt. Clin Microbiol Infect. 2013; 19(11):1082–8.

40. Touati M, Diene SM, Racherache A, Dekhil M, Djahoudi A, Rolain JM. Emergence of blaOXA-23 and blaOXA-58 carbapenemase-encoding genes in multidrug-resistant *Acinetobacter baumannii* isolates from University Hospital of Annaba, Algeria. Int J Antimicrob Agents. 2012; 40(1):89–91.

41. Abbott I, Cerqueira GM, Bhuiyan S, Peleg AY. Carbapenem resistance in *Acinetobacter baumannii*: laboratory challenges, mechanistic insights and therapeutic strategies. Expert Rev Anti Infect Ther. 2013; 11:395–409.

42. Mohamed NM, Raafat D. Phenotypic and genotypic detection of metallo-beta-lactamases in imipenem-resistant *Acinetobacter baumannii* isolated from a tertiary hospital in Alexandria, Egypt. Res J Microbiol. 2011; 6:750–60.

43. Espinal P, Poirel L, Carmeli Y, Kaase M, Pal T, Nordmann P, Vila J. Spread of NDM-2-producing *Acinetobacter baumannii* in the Middle East. J Antimicrob Chemother. 2013; 68:1928–30.

